# Comparative genomics and microbiome profiling reveal a conserved Vibrio rich mucus microbiota of the Florida false coral (*Ricordea florida*)

**DOI:** 10.64898/2026.09.17.752451

**Authors:** Megan E. Maloney, Eric C. Olson, Anthony G. Moss, Mark R. Liles, Nanette E. Chadwick, Katherine M. Buckley

## Abstract

Mucus is a critical interface between marine invertebrates and their environment and plays key roles in host defense. Despite its ecological importance, little is known about the microbial communities associated with mucus-producing corallimorpharians. Here, we used a multi-omics approach to characterize the mucus microbiome and immune repertoire of the Florida false coral, *Ricordea florida*. Culture-dependent and culture-independent analyses revealed that *R. florida* mucus harbors a bacterial community distinct from surrounding seawater and consistently dominated by *Vibrio* species. Whole-genome sequencing of 13 cultured *Vibrio* isolates representing nine distinct lineages revealed substantial taxonomic diversity, including several highly divergent strains that may represent undescribed species. Comparative genomic analyses identified enrichment of genes associated with carbohydrate acquisition, glycoside hydrolysis, and host colonization, including multiple components of the Tad/Flp adhesion system, suggesting adaptation to the mucus microenvironment. To investigate host factors that may shape microbial associations, we assembled and annotated a host transcriptome from healthy and immune-challenged polyps. This analysis identified a diverse innate immune repertoire, including Toll-like receptors, NOD-like receptors, lectins, scavenger receptors, complement-associated proteins, antiviral defense pathways, and membrane attack complex/perforin domain-containing effectors. Together, these findings demonstrate that *R. florida* supports a conserved, *Vibrio*-rich mucus microbiome and possesses a complex innate immune system capable of mediating host-microbe interactions at the mucosal surface. This study provides a foundation for understanding microbial colonization, immune defense, and holobiont function in an understudied cnidarian lineage.

## Introduction

Changing global climates and ecosystem loss due to human disturbance have deeply impacted marine invertebrate life and led to devastation of coral reef environments. As adults, most cnidarians are sessile and cannot relocate to overcome shifting environmental conditions but instead rely on specific protective phenotypes. One of these is the production and release of mucus, which has been implicated in defenses against environmental stress and predators as well as protection from microbial challenges [1]. Mucus contributes to the first line of defense by acting as a physical barrier that limits microbial entry. Molecularly, mucus is a secreted substance consisting of water, inorganic salts, carbohydrates lipids and glycoproteins, particularly mucins [2].

Many cnidarians rely on mutualistic relationships with algal symbionts, some of which contribute to mucus production [3–6]. Disruption of this relationship during bleaching can alter mucus secretion and composition, leading to shifts in associated microbial communities [6–9]. In reef-building corals, mucus-associated bacteria may also contribute to host defense [8, 10]. Disruption of the prokaryotic communities and subsequent mucus niche has been associated with decreased coral health [11]. These observations suggest that mucus microbiomes may play important roles in host health and resilience. Whether mucus-associated microbiomes perform similar functions in non-scleractinian cnidarians remains poorly understood. A recent study of the upside-down jellyfish *Cassiopea xamachana* found that its mucus microbiome is dominated by Alpha- and Gammaproteobacteria, similar to patterns observed in other symbiotic cnidarians [6]. Soft-bodied cnidarians provide an opportunity to study the complex relationships between host and bacteria that occupy the mucus layer and how these relationships are shaped by the host immune response.

The Florida false coral *Ricordea florida* is a tropical corallimorpharian that inhabits the Caribbean Sea and secretes a thick layer of mucus that covers external surfaces [12]. Specific functions for this mucus have not been investigated, although it has been proposed to serve as a physical barrier that protects the animal. The mucus also provides a nutrient-dense environment that is likely to support microbial life, although little is known about the bacterial species that inhabit the mucosal layer of corallimorpharians.

Here, we used a multi-omics approach to characterize the bacterial assemblage of *R. florida* mucus as well as the transcriptional immune response deployed by the corallimorpharian host. Using a combination of culture-dependent and culture-independent strategies, we find that the corallimorph mucus is enriched in a variety of *Vibrio spp.* and maintains a distinct microbial community from the surrounding seawater. Whole-genome sequencing of 13 *Vibrio* isolates reveals genes encoding a variety of metabolic and potentially pathogenic effector molecules.

Finally, analysis of transcriptome data reveals that the corallimorph immune response is mediated by a complex immune system consisting of receptors, signaling molecules and effector genes. Together, this work highlights fundamental aspects of animal immunity in a mucosal context and will facilitate future studies that focus on environmental drivers of disease and cnidarian immune responses.

## Materials and Methods

### Animal collection and maintenance

Individuals of *R. florida* were collected from shallow, nearshore sites in the Florida Keys, USA in summer 2018. Animals were housed and clonally propagated at Auburn University in recirculating aquaria with artificial seawater (ASW; Instant Ocean; 35 ppt) at 25°C and fed Sally’s Emerald Entrée Frozen Fish Food pellets (San Francisco Bay Brand) weekly.

### Isolation of culturable bacterial strains

Mucus samples (1 mL) were collected from six adults by tilting animals over 1.5 mL centrifuge tubes and collecting the run-off. To ensure that the sample was mucus, collections began only once the sample viscosity increased (**Supplemental Figure 1**). Seawater samples were collected from the tanks at the same time. Mucus and seawater samples were serially diluted 1:10 three times in filter-sterilized (0.22 µm) ASW. Serial dilutions (1/10 volume; 100 µl total; dilutions 10^−2^, 10^−3^, and 10^−4^) were plated onto Marine Agar 2216 (Difco Laboratories) and incubated at ∼26 °C for 24 hours. This reflects the natural temperature range of *R. florida* [12]. Colony forming units (CFUs) were counted and photographed to document morphology. Three representatives of each unique colony morphology were streaked onto marine agar plates to isolate pure bacterial cultures, and cryopreserved (20% v/v glycerol) at −80 °C.

Bacterial isolates were identified by 16S rRNA gene sequencing. Isolates were cultured on marine agar for 1 week at ∼25 °C. The 16S rRNA genes were amplified directly from colonies using the universal bacteria primer set 27F [13] and 1492R [14] and the Go-Green™ PCR master mix (Lucigen Corp.). Amplified products were sequenced with Sanger chemistry by Eurofins Operon (Huntsville, AL). Sequence reads were quality-trimmed using the CLC Genomics Workbench (Qiagen Inc., Aarhus, Denmark).

Taxonomic identification of cultured isolates was based on full-length 16S rRNA gene sequences. Consensus sequences were quality-trimmed and compared against the SILVA SINA Search and Classify service and the NCBI 16S rRNA database using BLASTn [15, 16]. Taxonomic assignments were based on agreement between the highest-scoring matches from both approaches and were reported at the phylum, genus, and, where possible, species level (**Supplemental Table 1**).

Isolates representing unique *Vibrio* ribotypes were selected for whole-genome sequencing. Bacterial strains were cultured on marine agar plates for 7 days. Genomic DNA was isolated using the E.Z.N.A. Bacterial DNA kit (Omega Biotek, Atlanta, GA), quantified using a Qubit fluorometer (ThermoFisher, Waltham, MA) and subjected to high-throughput sequencing using an Illumina NovaSeq 6000 (University of Illinois-Chicago’s Sequencing Core). Raw sequence reads were quality-trimmed (Q score >30) and assembled *de novo* using the CLC Genomics Workbench, v12.0 (Qiagen). Some isolates were re-sequenced using the Plasmidsaurus Whole Genome Sequencing service (Oxford Nanopore, R10.4.1 long-reads and *de novo* genome assembly). Genome details are shown in (**Supplemental Table 2**).

### Culture-independent analysis of bacterial communities

Mucus and seawater samples were collected as described above. From each of three separate aquaria, mucus was collected from two genetically distinct animals. DNA was isolated using the PowerWater kit (MO BIO Laboratories, Carlsbad, CA) according to manufacturer protocols and quantified using a Nanodrop spectrophotometer (ThermoFisher).

The V4 region of the 16S rRNA gene was amplified using barcoded primers and sequenced on an Illumina MiSeq platform at the University of Illinois-Chicago Sequencing Core. Sequences were processed in R using DADA2 [17]. including quality filtering, denoising, read merging, ASV inference, and chimera removal. Taxonomy was assigned using the SILVA database (v138.2), and chloroplast and mitochondrial sequences were excluded. Alpha-diversity metrics (observed richness, Shannon diversity, and Simpson diversity) were calculated using phyloseq [18]**(Supplemental Table 3)**.

### Orthology Identification and Phylogenomic Inference

To identify orthologous genes and predict functional enrichments associated with mucus-dwelling *Vibrio*, we analyzed the nine isolates described above alongside 40 reference-quality genomes from NCBI (36 *Vibrio* strains and four outgroup taxa; **Supplemental Table 4**). Predicted protein sequences were clustered into orthogroups using OrthoFinder v3.1.4 [19] with default settings.

Single-copy orthologs were selected that encoded encoding the 26 longest protein sequences were selected for analysis. Protein sequences were aligned using MAFFT (v7.525) [20], trimmed using TrimAl (v1.5.1) under the - automated1 heuristic and concatenated into a supermatrix using AMAS.py [21]. Maximum likelihood phylogenetic trees were reconstructed using IQ-TREE 3 [22] with ModelFinder (-m MFP) to determine the best-fit substitution model for each partition (Version 2.1.3; [23]); branch support was assessed using 1,000 ultra-fast bootstrap replicates (-bb 1000).

Carbohydrate-active enzymes (CAZymes) were annotated using dbCAN3 [24], retaining only predictions supported by at least two annotation methods. CAZyme annotations were mapped to orthogroups to examine the evolutionary distribution of carbohydrate-utilization functions across genomes. Differences in CAZyme family abundances among lineages were evaluated using Fisher’s exact tests with Benjamini-Hochberg correction (adjusted *P* < 0.05).

### Ecoplates

Microbial carbon utilization was assessed using Biolog EcoPlates as previously described [25]. Briefly, 150 μL of isolate A2 (10^5^ cells/mL), tank water, or mucus samples were inoculated into EcoPlates and incubated at 25 °C for 7 d. Optical density at 590 and 750 nm was measured at inoculation and after incubation, and carbon sources were considered positive when the change in optical density exceeded 1.0.

### Immune challenge and transcriptome analysis

Clonal *R. florida* individuals were generated via asexual budding and maintained on plexiglass tiles in aquaria. Two clonal polyps were isolated and placed in 500 mL beakers containing 300 mL ASW. *Vibrio* strain A2 (**Supplemental Table 2**), was grown overnight in marine broth. Bacteria were washed three times in filter-sterilized ASW (0.22 µm). One individual was lacerated using a sterile razor blade and exposed to 10^6^ bacteria/mL for 24 hours; the other polyp served as a non-injured, non-exposed control. Bacteria was added directly to polyps by pipetting 200 µL of washed bacteria directly over the oral cavity (**Supplemental Figure 1E,F**). Whole animals were collected 24 hr after exposure.

### Sequencing and transcriptome assembly

Total RNA was extracted using TRIzol (Invitrogen, Life Technologies), cleaned (Zymo RNA clean up) and quantified (Qubit 2.0 fluorometer, Life Technologies). Equal amounts of RNA from control and immune-challenged polyps were pooled and submitted to Novogene for poly(A)-selected mRNA library preparation and Illumina MiSeq sequencing (150 bp paired-end reads

Bioinformatic analyses were performed using the Alabama Supercomputer Authority. Raw reads were assessed with FastQC and quality filtered using TrimGalore v0.6.10 [26]. A *de novo* transcriptome assembly was constructed using Trinity v2.11.0 (--min_contig_length 300; [27]). To remove reads derived from algal symbionts, trimmed reads were mapped against a single symbiont index consisting of genome sequences from *Symbiodinium microadriaticum* strain:CCMP246 (GCA_001939145.1)*, Cladocopium goreaui* strain:SCF055 (GCA_947184155.2), and *Breviolum minutum* strain:Mf 1.05b.01 (GCA_000507305.1) [28]. Paired-end reads that failed to align concordantly were retained and used for downstream analyses.

A *de novo* transcriptome assembly generated 270,623 transcripts. Following removal of transcripts shorter than 250 bp and low-abundance transcripts (TPM < 1, quantified with Salmon Quant v1.10.0; [29]), 125,577 sequences were retained. Coding regions predicted with TransDecoder (v5.5.0) yielded 57,104 coding sequences (CDSs), of which 32,263 received functional annotations through EggNOG (v2.1.13; [30]). Transcripts assigned to Eukaryota, Opisthokonta, or Metazoa were retained and clustered at 95% sequence identity using CD-HIT (v4.8.1; [31]), producing a final reference transcriptome of 21,214 annotated sequences.

Transcriptome completeness was assessed using BUSCO (v.5.5.0; [32]).

## Results

### Corallimorph mucus is enriched in microbes relative to the surrounding seawater

To determine if corallimorph mucus harbors a bacterial community that is distinct from the environment, mucus and seawater samples were collected and plated on marine agar and incubated at 26 °C. This temperature reflects the typical *in situ* temperatures for *R. florida*, thereby increasing the likelihood of detecting ecologically relevant culturable taxa. Results indicate that plates inoculated with *R. florida* mucus exhibit consistently higher CFU counts than those that harbored seawater (on average, mucus samples contained 2.8x more CFU than seawater samples; **Figure 1A, Supplemental Figure 1B,C**). Throughout the course of this work, we repeated similar experiments several times using different growth temperatures, incubation times, dilution factors and a variety of corallimorph genetic backgrounds and consistently observed that plates inoculated with mucus contained more colonies than their seawater counterparts. Notably, colonies isolated from mucus also exhibited substantial phenotypic diversity (**Supplemental Figure 1D**). These isolates ranging from deeply pigmented colonies to cream, white, and peach morphotypes. One isolate exhibited an unusual agar-pitting phenotype, pointing to potential agarase activity which is often associated with specialized carbohydrate-degrading marine bacteria [33]. Together, these results suggest that *R. florida* mucus supports a diverse bacterial community that is distinct from that of the surrounding seawater.

**Figure 1:**
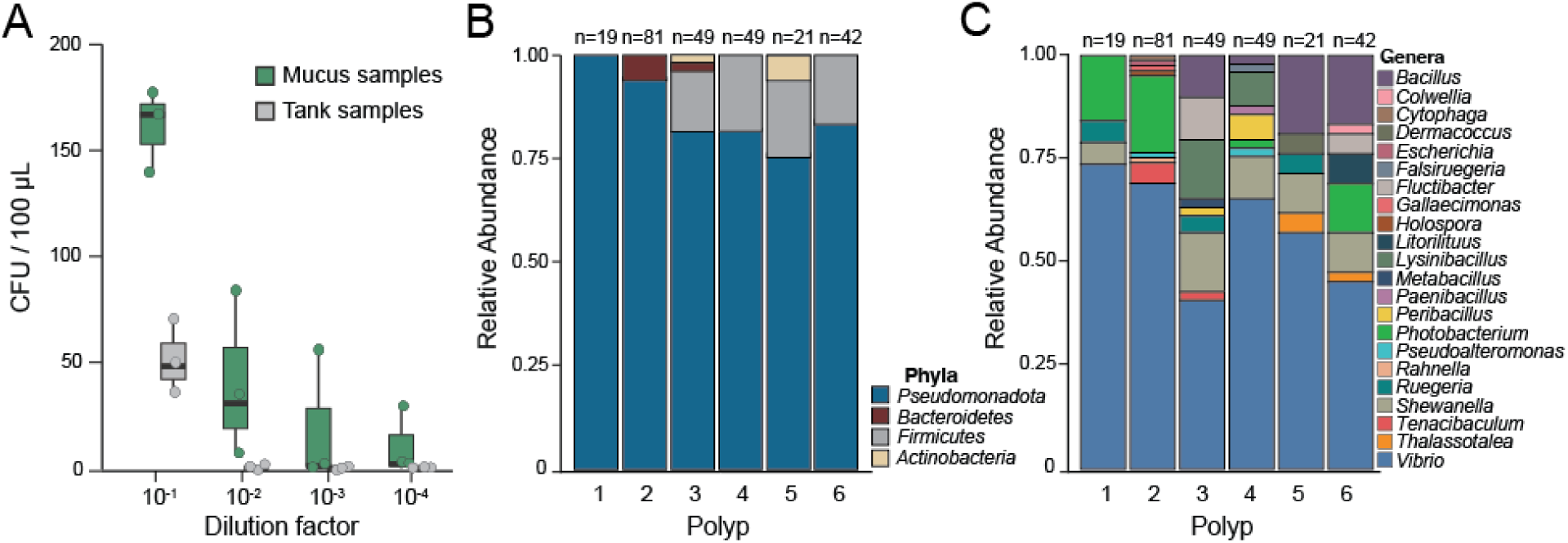
Corallimorph mucus is enriched for culturable bacteria relative to tank water and supports a diverse bacterial assemblage. **A. Culturable bacteria were consistently more abundant in *R. florida* mucus than in surrounding seawater.** Colony-forming unit (CFU) counts per plate are shown for mucus (green) and tank water (gray). Each sample was plated in triplicate, and individual plate counts are displayed. B,C. The culturable bacteria isolated from corallimorph mucus is dominated by members of the phylum *Pseudomonadota*. Taxonomic classification of the 261 culturable isolates recovered from mucus samples collected from six individual polyps revealed that, at the phylum level (B), isolates were overwhelmingly identified as *Pseudomonadota*. At the genus level (C), isolates were dominated by *Vibrio* spp., with additional representation from genera commonly found in marine environments. Similar taxonomic patterns were observed among the six polyps.

### Culturable bacteria within corallimorph mucus are enriched in Vibrio species

To determine whether the observed phenotypic heterogeneity reflected the presence of multiple distinct bacterial strains within corallimorph mucus, mucus samples were collected from six individual *R. florida* polyps housed within a single aquarium. Microorganisms were cultured as described above; colonies exhibiting distinct morphological characteristics were selected for further isolation and analysis. Through this approach, a total of 261 pure bacterial cultures were obtained. The complete 16S rRNA gene was amplified and sequenced from each isolated colony and the resulting sequences were analyzed using SILVA and BLAST to assign taxonomic identity at the phylum and genus levels (**Figure 1B,C**; **Supplemental Table 1**). Similar taxonomic patterns were observed across all six polyps, suggesting that individual variation was not a major contributor to the cultured mucus microbiome.

The culturable bacterial community isolated from *R. florida* mucus was dominated by *Pseudomonadota* (formerly *Proteobacteria*), which accounted for 207 of the 261 isolates (79.3%; **Figure 1B,C**). This taxonomic composition is consistent with bacterial communities commonly observed in marine environments [34, 35]. Furthermore, the majority of isolates could be assigned as *Vibrio sp.*, with 153 strains (58.6% of the total) assigned to this genus (**Figure 1C**). Among the culturable *Vibrio* species identified, *Vibrio coralliilyticus* (14 isolates) and *Vibrio hepatarius* (12 isolates) were the most frequently recovered. *V. coralliilyticus* is a well-characterized pathogen of marine corals and oysters [36], although no visible disease symptoms were observed in the animals sampled in this study. *V. hepatarius* is an aerobic, mesophilic bacteria that was originally isolated from the hepatopancreas of the shrimp *Littopanaeus vannamei* [37].

In addition to *Vibrio*, culturable bacteria isolated from *R. florida* mucus included representatives of the genera *Shewanella* (20 isolates), *Bacillus* (20 isolates), and *Photobacterium* (17 isolates) (**Figure 1C**). *Shewanella maritima*, the most frequently isolated *Shewanella* species, is a facultative anaerobe originally described from marine environments [38]. *Bacillus cereus* (eight isolates) is frequently associated with terrestrial and soil habitats but has also been recovered from marine and deep-sea environments [39]. Finally, members of the genus *Photobacterium* are closely related to *Vibrio* spp. [40] and are broadly distributed across marine ecosystems worldwide [41]. Together, analysis of the cultured bacterial assemblage indicates that corallimorph mucus is strongly enriched in *Vibrio* species while also supporting a diverse set of bacterial taxa commonly found in marine environments or in close association with marine organisms.

### Culture-independent methods indicate that corallimorph mucus harbors a diverse bacterial community

One limitation of culture-based approaches is that they selectively recover microorganisms capable of growing under a narrow set of laboratory conditions (here, marine agar cultured at 26 °C). To obtain a more comprehensive characterization of the bacterial community associated with *R. florida* mucus, we performed 16S rRNA gene amplicon sequencing on mucus samples collected from six polyps representing three independent aquaria (tanks 12, 14, and 15), together with corresponding seawater samples. Although seawater samples were processed in parallel, sequencing depth was substantially lower than that obtained for mucus samples and was insufficient for robust downstream community analyses; therefore, only mucus samples were included in subsequent analyses. Sequencing depth was consistent among mucus samples (30,718 - 45,373 reads per sample). Following quality filtering and contaminant removal, the final dataset contained 185 bacterial ASVs across six samples. Alpha diversity was moderate and relatively consistent among mucus samples, with observed richness ranging from 59 to 76 ASVs, Shannon diversity values from 2.21 to 2.62, and Simpson diversity values from 0.77 to 0.86 (**Supplemental Table 2**).

Taxonomic profiling revealed remarkable consistency across samples, with *Pseudomonadota* accounting for 99.3-99.9% of all sequences recovered from the mucus microbiome. Within this assemblage, *Vibrio* was the dominant bacterial genus in every sample, representing 72.6-87.1% of all reads (**Figure 2A**). Other genera were detected at substantially lower relative abundances. The agreement between culture-dependent and amplicon-based analyses supports the interpretation that *Vibrio* species represent core members of the *R. florida* mucus microbiome.

**Figure 2.**
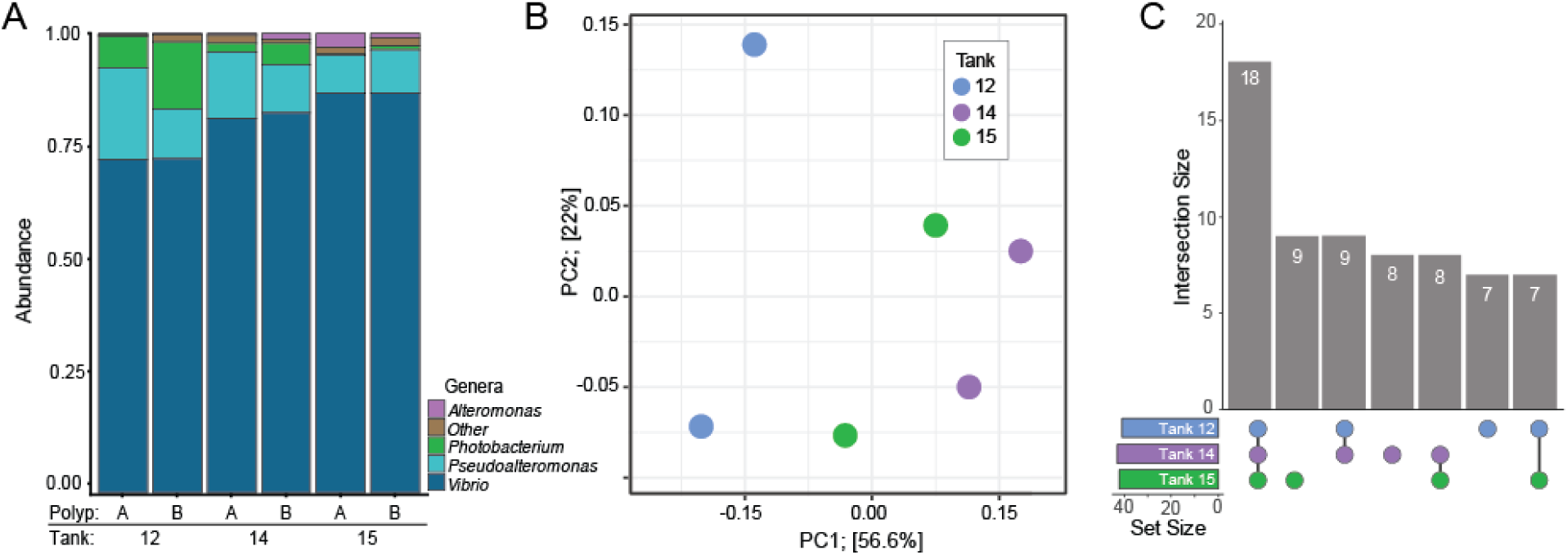
Culture-independent methods demonstrate that the microbial communities that inhabit Corallimorph mucus are enriched in *Vibrio* species. **A.** Relative abundance (%) of bacteria Genera collected from polyps from distinct tank systems reveals a dominance of *Vibrio*. Columns of each bar graph represent a polyp (A or B) and tank collected from is noted with the number. **B. Polyps from the same tank show no difference in bacterial assemblage.** A principal component analysis (PCA) reveals no differences in structure of bacterial communities between tank systems (12, 14, and 15 represented by blue, green and purple circles). X and Y axes show Principal Component 1 and Principal Component 2 that explain 56% and 22% of the total variance, respectively. **(C). Shared and unique ASVs among corallimorph-associated bacterial communities.** UpSet plot showing the overlap of amplicon sequence variants (ASVs) among Tanks 12, 14, and 15. An ASV was considered present within a tank only if it was detected in both animals sampled from that tank. Numbers indicate ASVs unique to individual tanks or shared among multiple tanks.

To assess variation in bacterial community composition among aquaria, Bray-Curtis dissimilarities were calculated from relative abundance data generated from the six mucus samples. Tank of origin explained approximately 39% of the observed variation in community composition (PERMANOVA; R² = 0.39, F = 2.56), although this effect was not statistically significant (p = 0.10). Samples collected from animals within the same aquarium generally exhibited lower Bray-Curtis dissimilarities than samples from different aquaria, suggesting modest aquarium-associated structuring of the mucus microbiome (**Figure 2B**). To compare bacterial taxa among aquaria, ASVs were filtered to retain only those detected in both animals sampled from a given tank before overlap analysis (**Figure 2C**). Of the 66 ASVs that met this criterion, 18 were shared among all three aquaria, and 24 were detected in at least two tanks. Together, these results indicate that although aquarium-associated variation was evident, many bacterial taxa were consistently associated with *R. florida* across tanks, suggesting the presence of a conserved mucus microbiota.

To assess whether compositional differences between mucus-associated and water-associated microbial communities were accompanied by differences in metabolic potential, we evaluated carbon substrate utilization using Biolog EcoPlates. Mucus-associated communities exhibited distinct substrate utilization profiles relative to surrounding tank water, including broader utilization of amino acids, carbohydrates, carboxylic acids, and phosphate-containing compounds (**Supplemental Figure 2**). These results provide additional evidence that the corallimorph mucus layer supports a functionally distinct microbial assemblage rather than a simple subset of the surrounding seawater community.

Analysis of both the culture-dependent and culture-independent strategies revealed that *Vibrio* spp. are the most prominent bacterial lineage associated with *R. florida* mucus. *Vibrio* represented the most abundant bacterial genus in every sample and was consistently detected across all sampled aquaria. The close agreement between culture-dependent and culture-independent approaches further suggests that these bacteria are well adapted to life within the corallimorph mucus layer. To investigate the genomic traits that may facilitate this association, representative *Vibrio* isolates were selected for whole-genome sequencing and comparative genomic analysis.

### Comparative genomic analyses identify genetic signatures of mucus association in Vibrio isolates

To gain a more comprehensive understanding of the *Vibrio* species associated with corallimorph mucus, 13 culturable isolates were selected for whole-genome sequencing (**Table 1, Supplemental Table 3**). These strains were chosen based on distinct colony morphologies and variation in 16S rRNA gene sequences. Assembled genome sizes ranged from 4.2 to 6.9 Mb and had an average GC content of 45.32%. These genomic characteristics are consistent with those reported for other members of the genus *Vibrio* [42].

**Table 1:** Gene content of *Vibrio* isolates.

| Isolate | Toxins | Flagella | Chemotaxis | Siderophore | Fe transport | Hemin | Hemolysin | Pili | Antimicrobial | Antibiotic | Secretion System |
| --- | --- | --- | --- | --- | --- | --- | --- | --- | --- | --- | --- |
| A2<br>(*V. owensii*) | 12 | 71 | 42 | 4 | 10 | 2 | 10 | 40 | 10 | 9 | 35<br>(II, IV, VI) |
| A4<br>(*V. coralliilyticus*) | 6 | 87 | 61 | 7 | 8 | 2 | 9 | 33 | 9 | 15 | 36,<br>(I, II, III, VI) |
| A6 | 6 | 46 | 45 | 3 | 9 | 2 | 9 | 26 | 9 | 10 | 37,<br>(I, II, VI) |
| C94 | 12 | 40 | 65 | 5 | 8 | 2 | 11 | 48 | 10 | 9 | 33,<br>(II, III, VI, VIII) |
| C130<br>(*V. paucivorans*) | 10 | 44 | 58 | 4 | 7 | 2 | 11 | 50 | 7 | 7 | 33,<br>(II, III, VI) |
| C136<br>(*V. mediterranei*) | 16 | 86 | 38 | 3 | 9 | 2 | 8 | 48 | 11 | 8 | 40,<br>(I, II, IV, VI, VIII) |
| C147<br>(*V. ishigakensis*) | 6 | 2 | 1 | 1 | 3 | 2 | 5 | 44 | 8 | 7 | 30<br>(II, VI) |
| D58 | 12 | 43 | 76 | 2 | 6 | 2 | 6 | 36 | 8 | 6 | 35<br>(II, III, VI, VIII) |
| D75 | 13 | 71 | 75 | 2 | 3 | 2 | 6 | 38 | 8 | 4 | 18,<br>(II, IV, VI, VIII) |
| *V. harveyi* | 16 | 84 | 42 | 8 | 1 | 0 | 2 | 47 | 0 | 0 | 147,<br>(I, II, III, IV, VI) |
| *V. diazotrophicus* | 5 | 80 | 53 | 8 | 0 | 1 | 3 | 7 | 0 | 3 | 50,<br>(I, II, IVB, VI) |
| *V. cholerae* | 28 | 46 | 58 | 4 | 1 | 1 | 3 | 8 | 0 | 3 | 57,<br>(I, II, III, IVB, VI) |

Pairwise average nucleotide identity (ANI) analyses were performed to identify closely-related isolates and assess taxonomic diversity across the 13 *Vibrio* genome assemblies. This analysis identified three groups of isolates with >95% identity: A4/B77/C98, C130/D80, and D58/D75/G20 (**Figure 3A**). The remaining six isolates did not cluster with any other genomes in the dataset and were therefore treated as distinct species-level lineages. Species-level assignments were further evaluated using GTDB-Tk, MiGA, and TYGS. Four species could be confidently assigned to isolates, including *V. coralliilyticus* (the A4/B77/C98 cluster), *V. mediterranei* (C136), *V. owensii* (A2), and *V. paucivorans* (C130/D80). Notably, several of these species have previously been associated with coral disease, bleaching, or tissue loss [36, 43, 44]. Together, these results reveal substantial taxonomic diversity among mucus-associated *Vibrio* isolates, including both recognized coral-associated species and several deeply divergent lineages.

**Figure 3:**
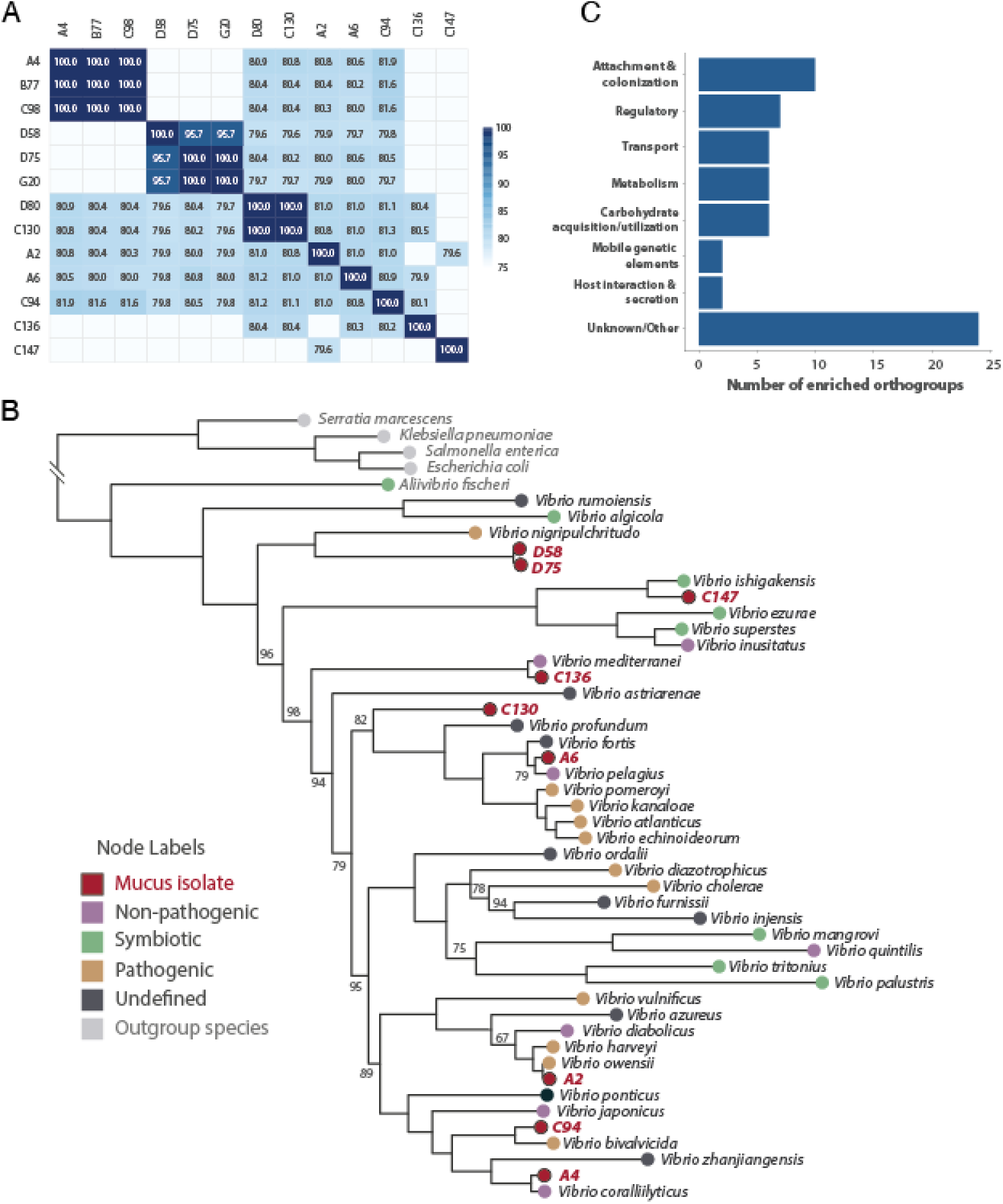
Phylogenetic analysis and functional analysis of *Vibrio* mucus isolates. **A. Corallimorph mucus harbors potentially novel *Vibrio* strains.** Average Nucleotide Identity (ANI) matrix comparing genomic similarity among isolated strains. Color scale indicates percentage identity (range: 75%–100%). White boxes indicate that the genomic similarity was less than 75%. **(B). *Vibrio* isolated from Corallimorph mucus span a wide diversity of clades and ecological niches.** Maximum-likelihood phylogenetic tree based on amino acid sequences from 26 single copy orthogroups showing the placement of mucus isolates (highlighted with red nodes and red text labels) relative to reference species across the *Vibrio* genus. Tip nodes are color-coded by ecological classification: mucus isolate (this study, red), non-pathogenic (purple), symbiotic (green), pathogenic (brown), undefined (dark grey), and outgroup species (light grey). All nodes are supported by 100% bootstrap support except where otherwise noted. **(C). Genes associated with attachment and colonization were the most enriched functional category among mucus-isolated *Vibrio* genomes.** Enrichment analysis showing the number of enriched orthogroups within each functional category. Twenty-three enriched orthogroups were grouped as unknown/other, which encompasses both poorly annotated genes and genes with functions not readily assigned to the major categories.

The remaining five isolates exhibited varying degrees of taxonomic uncertainty. Isolate C94 was classified as *V. tubiashii* by both MiGA and GTDB-Tk; however, its ANI value of 95.7% lies near the lower boundary typically used for species delineation, which is consistent with the possibility that this is a divergent lineage within *V. tubiashii*. Isolate A6 was classified as *V. fortis* A by GTDB-Tk but shared only 90.3% ANI with its closest MiGA reference genome (*V. pelagius*), indicating substantial divergence from currently recognized species. Similarly, isolate C147 shared only 84.4% ANI with *V. ishigakensis* and could not be assigned to a described species by either GTDB-Tk or TYGS. The three closely related isolates D58, D75, and G20 exhibited average pairwise ANI values of 98.1% but shared only ∼75.7% ANI with their closest reference genome (*V. penaeicida*), indicating that they represent a single, deeply divergent lineage. Representative genomes from these groups were compared against all available *Vibrio* type strain genomes using the Type Strain Genome Server (TYGS). Digital DNA-DNA hybridization (dDDH) values were well below the 70% species threshold for all reference strains, providing strong support that C147 and the D58/D75/G20 lineage represent previously undescribed *Vibrio* species. Because D58, D75, and G20 were nearly identical at the genome level, subsequent analyses were performed using D75 as the representative genome for this lineage.

To examine the evolutionary relationships of the mucus-associated *Vibrio* isolates, we reconstructed a genome-scale phylogeny using 26 single-copy orthologs identified across all analyzed genomes and a reference dataset representing diverse *Vibrio* species and their closest relatives identified by FastANI (Supplemental Table 4). Protein sequences were aligned, concatenated, and used to infer a phylogeny that largely corroborated species assignments based on genomic similarity analyses (Figure 3B). Isolates A2, A4, C130, and C136 grouped with their respective reference species, while C94 clustered within the *V. tubiashii* lineage despite its borderline assignment by MiGA and GTDB-Tk. In contrast, A6 formed a sister lineage to *V. pelagius*, C147 formed a divergent sister lineage to *V. ishigakensis*, and D75/D80 formed a distinct lineage related to the *V. pelagius-V. nigripulchritudo* clade. The phylogenetic distinctiveness of C147 and the D75/D80 lineage supports their potential status as previously undescribed taxa. This phylogeny provides a framework for subsequent analyses of gene content and functional evolution.

Functional annotation revealed variable distributions of virulence-associated factors, adhesion machinery, antibiotic resistance determinants, and secretion systems across the mucus-associated *Vibrio* isolates. Given the pathogenic potential of many *Vibrio* species, we compared these profiles with those of two well-characterized pathogens, *Vibrio cholerae* and *Vibrio harveyi* (reference pathogenic vibrios; RPV). With the exception of *V. mediterranei* C136, which encoded 16 toxin-associated genes, mucus-associated isolates generally possessed fewer toxin-related genes than the RPV genomes, including RTX toxins, RTX-associated Ca^2+^-binding proteins, *zot*, and toxin-antitoxin system components (Table 1; Supplemental Table 5). Gene content related to antibiotic resistance also differed between groups. *V. corallilyticus* A4 and *V. pelagius* A6 encoded the largest numbers of resistance-associated genes (15 and 10, respectively), whereas no such matches were detected in the RPV genomes under the annotation scheme employed here. Secretion system repertoires likewise varied substantially; RPV genomes encoded a broader diversity of systems (Types I, II, III, IV, IVB, and VI) and a greater number of associated genes (50-147), whereas mucus-associated isolates typically encoded fewer secretion-related genes (30-40) and consistently possessed only Types II and VI. Together, these results indicate that mucus-associated *Vibrio* differ from reference pathogens in their complements of toxin, resistance, and secretion-system genes, suggesting distinct ecological strategies and host interactions.

To identify genomic features associated with mucus colonization, genome sequences from the nine taxonomically distinct strains were functionally annotated and clustered into orthologous gene groups. Enrichment analysis comparing the mucus-associated isolates to a phylogenetically diverse reference set of 40 *Vibrio* species identified 63 orthogroups that were significantly overrepresented in the mucus-associated isolates (adjusted p-value < 0.05). Based on functional annotation, these orthogroups were assigned to eight categories (**Supplemental Table 6**). Notably, orthogroups involved in carbohydrate acquisition and utilization as well as attachment and colonization were among the most abundant characterized functional categories, suggesting an enhanced capacity to exploit mucus-derived nutrients and establish stable associations with host mucus. Nearly half of the enriched orthogroups (26/63) lacked informative functional annotations, suggesting that potentially important mechanisms underlying mucus association remain poorly characterized.

The mucus-associated isolates were enriched in genes associated with carbohydrate uptake, utilization, and degradation relative to the reference *Vibrio* dataset (**Supplemental Table 6)**. These included a phosphotransferase system (PTS) IIB component (COG1264), which mediates carbohydrate transport and phosphorylation [45], and a β-fructosidase/levanase (COG1621) involved in the hydrolysis of fructose-containing polysaccharides [46]. Additional enriched orthogroups encoded a glycogen debranching enzyme glucanotransferase domain, a glycosyl hydrolase family 65 (GH65) domain-containing protein, and a member of glycosyl hydrolase family 32 (GH32), all of which are associated with the breakdown and utilization of complex carbohydrates [47–49]. To further evaluate genomic adaptations for carbohydrate utilization, we performed a comparative CAZyme analysis [50] of the mucus-associated isolates and the reference *Vibrio* genomes. Consistent with the orthogroup enrichment analysis, mucus-associated isolates exhibited an increased abundance of glycoside hydrolase families, including GH65 and GH13_4, which are involved in the degradation of complex α-glucans and related carbohydrates [51, 52]. Collectively, these findings point to an enhanced capacity to access, degrade, and utilize complex carbohydrates and support the hypothesis that these isolates have adapted to exploit the diverse glycans and polysaccharides present within the coral mucus microenvironment.

Orthogroups associated with attachment and colonization were also significantly enriched in the mucus-associated isolates (**Supplemental Table 6**). Notably, many of these genes encoded components of the Tad (tight adherence) locus and the Flp (fimbrial low-molecular-weight protein) pilus assembly system, including *tadG*, *tadD*, *cpaB*, *cpaE*, *tadE*-like proteins, and multiple Flp pilus structural and assembly proteins. The Tad/Flp system mediates the production of adhesive pili that facilitate surface attachment, biofilm formation, and stable colonization of host-associated environments. The enrichment of multiple components spanning pilus biogenesis, assembly, and secretion suggests that mucus-associated isolates possess an enhanced capacity to adhere to and persist within the coral mucus layer [53]. These findings indicate that successful exploitation of mucus-derived nutrients may be coupled with mechanisms that promote stable association with host surfaces.

The remaining enriched orthogroups spanned several functional categories, including regulation, metabolite transport, host interaction and secretion, core metabolism, and mobile genetic elements. Several transcriptional regulators and signaling proteins suggest an enhanced capacity to respond to fluctuating environmental conditions within the mucus microenvironment. Enriched transport systems may facilitate the uptake of host-derived nutrients, while mobile genetic elements indicate genome plasticity that could promote adaptation to host-associated lifestyles [54]. Finally, 26 enriched orthogroups lacked informative functional annotations, indicating that a substantial portion of the genetic basis underlying mucus association remains poorly understood.

### Corallimorphs express an array of transcripts with potential roles in immunity

Because host factors likely contribute to microbiome structure, we characterized the immune repertoire of *R. florida* using transcriptome sequencing. RNA from healthy and immune-challenged clonal polyps was pooled to maximize transcript diversity. Following removal of putative symbiont-derived sequences, *de novo* assembly and annotation yielded 21,214 non-redundant host transcripts with confident functional assignments (Table 1). These annotations were screened for genes associated with innate immunity, including pattern recognition receptors (PRRs), signaling molecules, and immune effector proteins.

Annotation of the *R. florida* transcriptome identified a diverse innate immune repertoire, including multiple classes of pattern-recognition receptors (PRRs), signaling molecules, and downstream effectors (**Table 2**). A complete TLR-signaling pathway was detected, including putative TLRs, MyD88-like adaptors, SARM-like proteins, and numerous TIR domain-containing proteins, as well as TRAF-family signaling molecules and regulators of NF-κB activity. Additional PRRs included extensive repertoires of lectin-containing and scavenger receptor cysteine-rich (SRCR) proteins, suggesting substantial capacity for recognition of diverse microbial and environmental ligands at the mucus-covered host surface.

**Table 2:** Diversity and abundance of innate immune gene families identified in the *R. floridae* transcriptome.

| Functional Category | Gene Family/Component | Number of transcripts |
| --- | --- | --- |
| **Pattern Recognition Receptors** | Toll-like receptors (TLR); LRR-TIR | 4 |
|  | Nod-like receptors (NLR); NACHT domain + effector domain | 51 |
|  | Scavenger receptor (SRCR) | 43 |
|  | Lectin-like PRRs | 41 |
| **Nucleic Acid Sensing and Antiviral Defense** | Rig-I like receptors (RIG-I/DDX58) | 3 |
|  | Helicase sensors (DDX41, DDX60, DHX9, DHX15, DHX33) | 12 |
|  | RNA editing (ADAR, ADARB1, ADARB2) | 6 |
|  | RNAi (DICER1) | 3 |
| **Immune signaling** | BCL10 | 1 |
|  | TNFAIP8 | 1 |
|  | TNFAIP3/A20 | 1 |
|  | CARD-DEATH protein | 1 |
|  | TRAF-domain proteins | 35 |
|  | MAPK signaling components | 19 |
| **Adaptor Proteins** | MyD88 (DEATH-TIR) | 1 |
|  | SARM (SAM- TIR) | 2 |
|  | TIR domain (TIR domain only) | 12 |
|  | MAVS (1) | 1 |
| **Transcriptional Regulators** | ATF | 2 |
|  | ETS | 12 |
|  | GATA | 2 |
|  | IRF | 2 |
|  | Rel-family transcription factors | 8 |
|  | BCL3 | 2 |
| **Cell death** | BCL proteins | 3 |
|  | Initiator Caspases (CARD-caspase) | 4 |
|  | APAF1(CARD,NB-ARC,WD40) | 4 |
|  | Executioner Caspases (caspase domain) | 2 |
|  | Atypical caspases (Ig-caspase) | 4 |
| **Complement related proteins** | C1q-like proteins | 8 |
|  | CSMD | 5 |
|  | CUB/Sushi (3) | 2 |
|  | Complement-associated lectins | 12 |
| **Cytokines and receptors** | MIF | 1 |
|  | TNFSF | 4 |
| **Effectors** | Cathepsin | 14 |
|  | MACPF | 31 |

Putative intracellular immune sensors were also abundant, including NACHT-containing proteins with CARD, PYRIN, LRR, and SPRY domains characteristic of NLR-like receptors. The transcriptome further contained multiple RNA-sensing helicases, RNA interference components, and RNA editing enzymes, including DDX-family helicases, RIG-I-like receptors, MAVS, DICER1, and ADAR-family proteins, consistent with conserved antiviral defense pathways. Several transcripts encoded proteins associated with complement-like recognition systems, including C1q-, CUB-, Sushi-, and lectin-containing proteins, as well as CSMD family members and complement receptor-like molecules. Components of the terminal complement pathway (C5-C9) were not identified. Predicted immune effector genes included cathepsins, caspases, BCL10, APAF1, and an expanded repertoire of membrane attack complex/perforin (MACPF) domain-containing proteins, indicating diverse mechanisms for immune signaling, apoptosis, and antimicrobial defense. The extensive repertoires of lectins, SRCR proteins, NLR-like receptors, and complement-associated proteins identified here may provide mechanisms through which *R. florida* discriminates among and regulates microbes residing within the mucus layer.

## Discussion

Here we present a multifaceted approach to characterize the microbial communities associated with the mucus layer of the corallimorpharian *R. florida* and reveal that this microbiome is distinct from the surrounding seawater and consistently dominated by *Vibrio* species. Comparative genomics identified diverse *Vibrio* lineages, including potentially undescribed taxa, and revealed enrichment of genes associated with carbohydrate utilization and host colonization. In contrast, host transcriptome annotation identified a diverse innate immune repertoire, including pattern-recognition receptors, immune signaling molecules, and effector proteins, suggesting that both microbial and host factors contribute to microbiome assembly.

Together, these findings provide new insights into host-microbe interactions at the mucosal surface of *R. florida* and establish a foundation for understanding microbiome assembly and immune function in corallimorpharians.

Vibrio species are widespread members of marine microbial communities and occur as free-living, commensal, mutualistic, and pathogenic associates of diverse marine organisms, including mollusks, fish, corals, and anemones [55, 56]. Several *Vibrio* species, including *V. shiloi* and *V. coralliilyticus*, are well-known coral pathogens associated with bleaching, tissue loss, and disease [57–59]. However, focusing exclusively on pathogenic interactions may overlook the broader ecological importance of this genus. Here, both culture-dependent and culture-independent approaches identified *Vibrio* as the dominant bacterial lineage associated with apparently healthy *R. florida* individuals, suggesting that these bacteria represent stable components of the corallimorph mucus microbiome rather than transient opportunists. The recovery of multiple phylogenetically diverse *Vibrio* species, including several potentially undescribed lineages, further suggests that the mucus habitat supports a diverse assemblage of bacteria that may occupy distinct ecological roles. Together, these findings support a growing view that many host-associated *Vibrio* species function as persistent members of marine microbiomes [60] and that pathogenicity represents only one outcome within a broad spectrum of possible host-microbe interactions.

Our findings suggest that corallimorph mucus represents a specialized ecological niche that contributes to host defense and shapes microbial community composition. In hexacorals, mucus contains a diverse mixture of host- and symbiont-derived organic compounds that create a nutrient-rich microenvironment [62, 63]. Consistent with this, comparative genomic analyses revealed that mucus-associated *Vibrio* isolates were enriched in genes involved in carbohydrate acquisition and utilization, including multiple glycoside hydrolases, suggesting that the ability to access host-derived glycans may contribute to successful colonization. Mucus-associated isolates were also enriched in components of the Tad/Flp adhesion system, which mediates surface attachment and biofilm formation [64–66]. Notably, we find that these traits were shared among multiple phylogenetically distinct *Vibrio* lineages. This pattern suggests that adaptation to the mucus environment is not restricted to a single evolutionary lineage but instead has arisen repeatedly among diverse *Vibrio* species. Such convergence is consistent with the hypothesis that the corallimorph mucus layer acts as a complex environmental filter that favors bacteria possessing traits required for nutrient acquisition and stable host association.

Transcriptome annotation revealed that *R. florida* possesses a diverse innate immune repertoire composed of pattern-recognition receptors, signaling molecules, transcriptional regulators, and putative effector proteins. Components of several canonical animal immune pathways were identified, including TLRs and MyD88-dependent signaling, NLR-like receptors, antiviral sensors, complement-associated proteins, and membrane attack complex/perforin (MACPF) domain-containing effectors. Many of these gene families have been reported previously in anthozoans and other cnidarians [1, 67–70], where they are have been implicated not only in pathogen defense but also the regulation of symbiotic and commensal microbes Expansions of innate immune receptor families are increasingly recognized as a hallmark of immunity in basal metazoans [67, 71–76]. Consistent with this pattern, the extensive repertoires of lectins, SRCR proteins, NLR-like receptors, and complement-associated proteins identified in *R. florida* suggest substantial capacity for immune-mediated regulation of microorganisms within the mucus layer. The identification of these pathways in a corallimorpharian extends evidence for complex innate immunity beyond reef-building corals and suggests that mechanisms governing host-microbe interactions may be conserved across Hexacorallia. Together, these findings support a model in which microbiome assembly is shaped by reciprocal interactions between host immunity and associated microbial communities.

Increasing evidence suggests that microbial communities are integral components of cnidarian biology, influencing host nutrition, immune function, disease susceptibility, and responses to environmental stress [77]. In reef-building corals, shifts in microbial community composition often accompany bleaching, disease outbreaks, and other stress responses [11, 78], whereas stable microbial associations have been proposed to contribute to host health through nutrient cycling, competitive exclusion of pathogens, and the production of antimicrobial compounds [79]. Although these interactions have been investigated extensively in scleractinian corals, comparatively little is known about the ecological roles of mucus-associated microbes in corallimorpharians. The consistent association between *R. florida* and a diverse assemblage of *Vibrio* species suggests that these bacteria may represent more than transient environmental colonists and could contribute to normal holobiont function. Whether these interactions are beneficial, neutral, or context dependent remains unclear; however, the genomic diversity observed among the mucus-associated *Vibrio* isolates suggests that different lineages may occupy distinct ecological roles within the mucus microenvironment. Some lineages may contribute to nutrient acquisition through the degradation of complex carbohydrates, whereas others may influence microbial community structure through competitive interactions or the production of secondary metabolites. Determining how these bacteria contribute to host physiology, microbiome stability, and environmental resilience represents an important direction for future research.

Future work should focus on determining the functional consequences of these host-microbe associations. Integrating controlled colonization experiments, microbiome manipulation, and gene-expression analyses will help resolve whether mucus-associated *Vibrio* species contribute to nutrient cycling, pathogen resistance, or other aspects of holobiont function. Such studies will provide a deeper understanding of how microbial communities influence the ecology and resilience of corallimorpharians and other cnidarians. More broadly, these findings suggest that the mucus layer of *R. florida* represents a structured ecological niche shaped by both microbial adaptations for colonization and host mechanisms for microbial recognition. The discovery of a conserved, *Vibrio*-dominated microbiome, together with evidence for both bacterial specialization and a complex host immune repertoire, highlights the potential importance of host-microbe interactions in corallimorph biology. As a tractable aquarium model that can be clonally propagated, *R. florida* may also provide a valuable system for investigating the mechanisms governing microbiome assembly, immune regulation, and environmental resilience in cnidarians.

## Data Availability Statement

All genome sequences generated during and during the current study have been deposited at NCBI under BioProject PRJNA1525680 and PRJNA1366596. The raw sequencing reads from *R. florida* have been deposited at NCBI (accession SRR40318881). Additional files, including custom scripts, 16S rRNA sequences from cultured bacterial isolates and transcriptome assemblies are available at https://github.com/mem0294/Maloneyetal2026_Corallimorph_mucus.

## Acknowledgements

The authors would like to thank Megan Roberts and Taylor Plunkett for their assistance with colony isolation. This work was supported by an NSF award to K.M.B. (2131297).

## Supplemental Figures

**Supplemental Figure 1:**
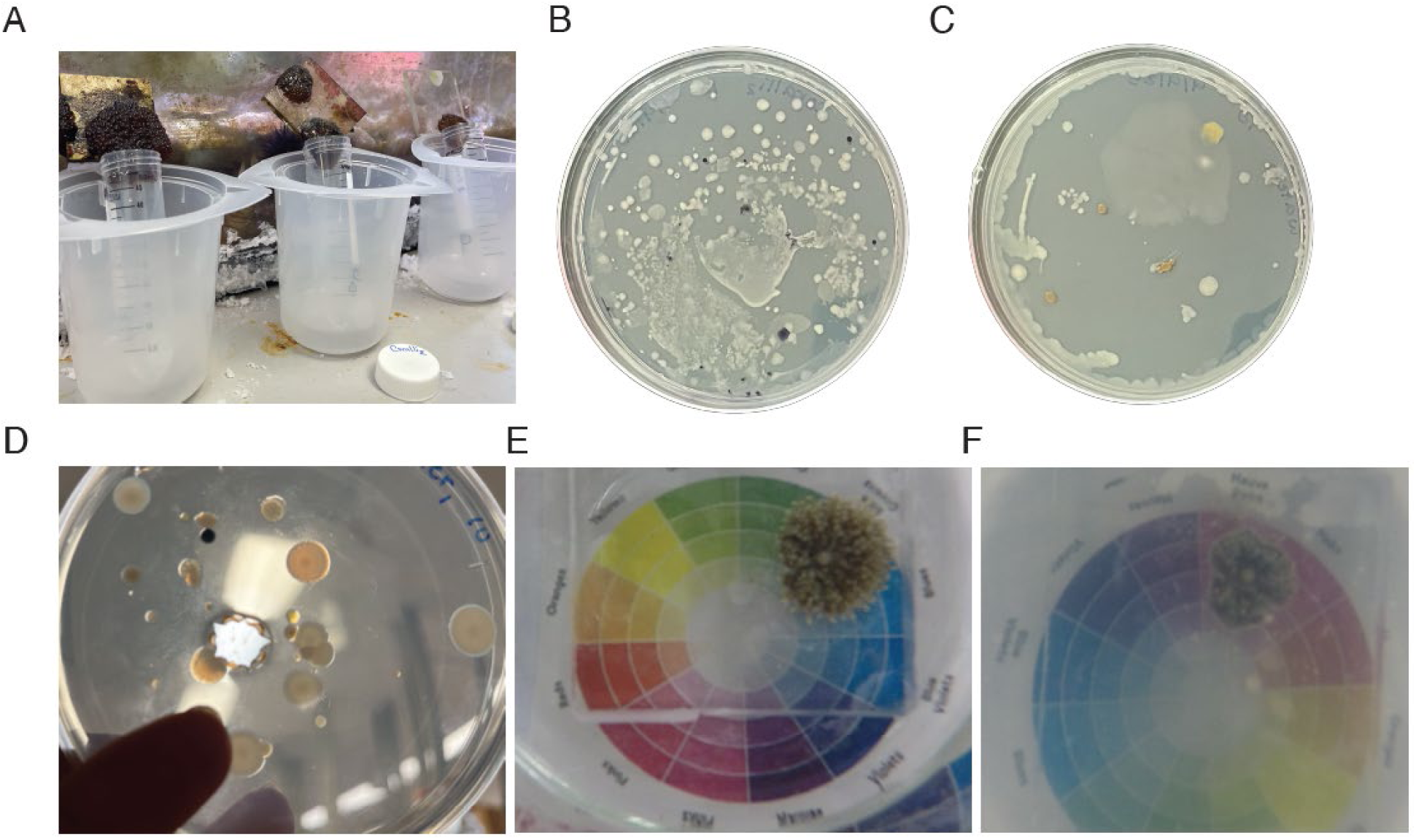
Mucus collection, bacterial culturing and *R. florida* infection methodology images. **(A) Corallimorpharian mucus collection.** After initial salt water dripped off animals and plates, mucus was collected from adult Corallimorpharian animals by tilting over 1.5 mL centrifuge tubes and collecting the run-off. **B,C) *R. florida* mucus exhibit consistently higher CFU counts than those that harbored seawater.** Corallimorph mucus (A) and surrounding sea water (C) were cultured on Marine Agar plates and the number of colony forming units were quantified. (**D) Corallimorpharian mucus harbors morphologically diverse bacterial isolates.** Colonies isolated from *R. florida* mucus exhibited substantial variation in pigmentation and morphology, including dark purple/black (A), translucent cream (B-D), white (E), and peach (F) morphotypes. Isolates also differed in colony architecture and growth characteristics. Among these, one isolate displayed an unusual agar-pitting phenotype (G), suggesting extracellular agar-degrading activity. (**E,F) *R. florida* were infected with A2 *Vibrio*.** Clonal polyps from control treatment (E) and high bacterial exposure (F) were pooled in equal quantities for mRNA library construction and sequencing using MiSeq technology (Illumina).

**Supplemental Figure 2:**
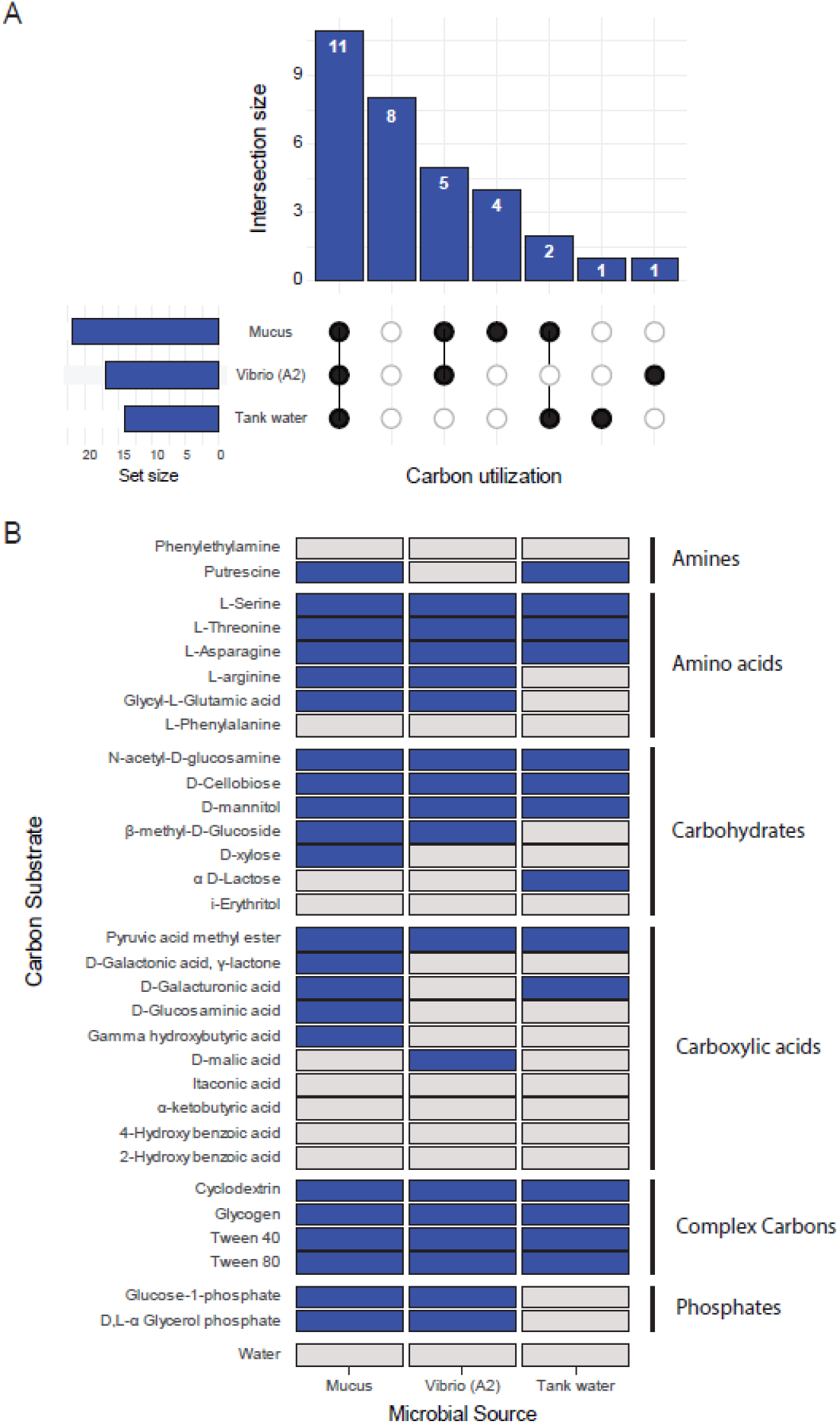
Carbon utilization profiles differ among mucus-associated microbial communities, tank water communities, and a representative mucus-associated *Vibrio* isolate. **(A)** UpSet plot showing the overlap in utilization of 31 carbon substrates among mucus samples, tank water samples, and *Vibrio* isolate A2. Bars indicate the number of substrates utilized by each unique combination of sample types. (**B**) Heatmap depicting utilization of 31 carbon substrates categorized into six functional groups: Amines, Amino Acids, Carbohydrates, Carboxylic Acids, Complex Carbon, and Phosphates. Separately, 150 μL of mucus, tank water, or *Vibrio* A2 cultures (10^5^ cells/ml) were inoculated into Biolog EcoPlates and incubated at 25°C. Optical density (OD) at 590 and 750 nm was measured at inoculation and after 7 days using a plate reader (Bio Stack Ready, USA). Blue boxes indicate positive substrate utilization (mean change in OD > 1.0), whereas light gray tiles indicate little or no detectable substrate utilization.

## Supplemental Tables

Supplemental Table 1: Characteristics of cultured bacterial isolates

Supplemental Table 2: Statistics of culture-independent 16S analysis

Supplemental Table 3: Vibrio genome stats

Supplemental Table 4: Phylogeny

Supplemental Table 5: Vibrio genome annotation results

Supplemental Table 6: Functional enrichments in mucus isolate genomes

Supplemental Table 7: *R. florida* immune transcriptome annotations

